# Decoding the Neural Dynamics of Headed Syntactic Structure Building

**DOI:** 10.1101/2024.11.07.622560

**Authors:** Junyuan Zhao, Ruimin Gao, Jonathan R. Brennan

## Abstract

The brain builds hierarchical phrases during language comprehension; however, the representational details and dynamics of the phrase-building process remain underspecified. This study directly probes whether the neural code of a phrase involves reactivating the syntactic property of a key subcomponent (the “phrasal head”). To this end, we train a part-of-speech sliding-window neural decoder (verb vs. adverb) on EEG signals recorded while participants (N = 30) read sentences in a controlled experiment. The decoder reaches above-chance performance that is spatiotemporally consistent and generalizes to unseen data across sentence positions. Appling the decoder to held-out data yields predicted activation levels for the verbal “head” of a verb phrase at a distant non-head word (adverb); the critical adverb appeared either at the end of a verb phrase or at a sequentially and lexically matched position with no verb phrase boundary. There is stronger verb activation beginning at ∼600 milliseconds at the critical adverb when it appears at a verb phrase boundary; this effect is not modulated by the strength of conceptual association between the two subcomponents in the verb phrase nor does it reflect word predictability. Time-locked analyses additionally reveal a negativity waveform component and increased beta-delta inter-trial phase coherence, both previously linked to linguistic composition, in a similar time window. With a novel application of neural decoding, our findings delineate the temporal dynamics by which the brain encodes phrasal representations by, in part, reactivating the representation of key subcomponents. We thus establish a link between cognitive accounts of phrase structure representations and electrophysiological dynamics.

## 1. Introduction

During language comprehension, the brain accesses the representation of words and combines them into phrases (Friederici et al., 2017; Hagoort, 2019; Pylkkänen, 2019). This process is structure-dependent in that its neural readout is modulated by hierarchically structured representations over and above sequential word inputs (Ding et al., 2016; Kaufeld et al., 2020; Nelson et al., 2017; Stanojević et al., 2023). Still, the detailed properties of the encoded representations remain subject to significant debate (c.f. Frank & Christiansen, 2018). One candidate drawn from linguistics is that the syntactic features of a phrase are derived from a nucleus word called the phrasal “head” (Chomsky, 1957; Jackendoff, 1977): A verb phrase, for example, is built around a verb. Such a component is necessary for any neural system that constructs recursive hierarchical linguistic structure (Sprouse & Hornstein, 2016). This theory carries the untested prediction that information about the head word can be recovered from a phrasal representation. We test this prediction with a novel two-step neural decoding pipeline on electroencephalography (EEG) to provide insight into (1) whether the neural activation of the head word is stronger at a temporally distal phrase closing time-point and (2) whether (re)activation of the head is modulated by conceptual information (Fedorenko et al., 2016; Parrish & Pylkkänen, 2022).

Linguistic theories hold that hierarchical structure building is contingent on endocentric phrase representations where a phrase shares syntactic properties with the “head” word (Bloomfield, 1933). This shared property is captured by the notion of part-of-speech (PoS): Phrases and words that share the same PoS can be exchanged with each other without changing grammaticality (Figure 1 and Carnie, 2021:78). Neural signals are robustly modulated by hierarchical syntactic structure beyond linear word encodings (Brennan et al., 2016; Coopmans et al., 2024; Kaufeld et al., 2020; Nelson et al., 2017) and oscillatory signatures reflect phrase boundaries independently of the sequential position of individual words (Burroughs et al., 2021; Lo et al., 2022; Zhao et al., 2024). Yet, these data do not bear on the neural representation of phrasal information.

**Figure 1.**
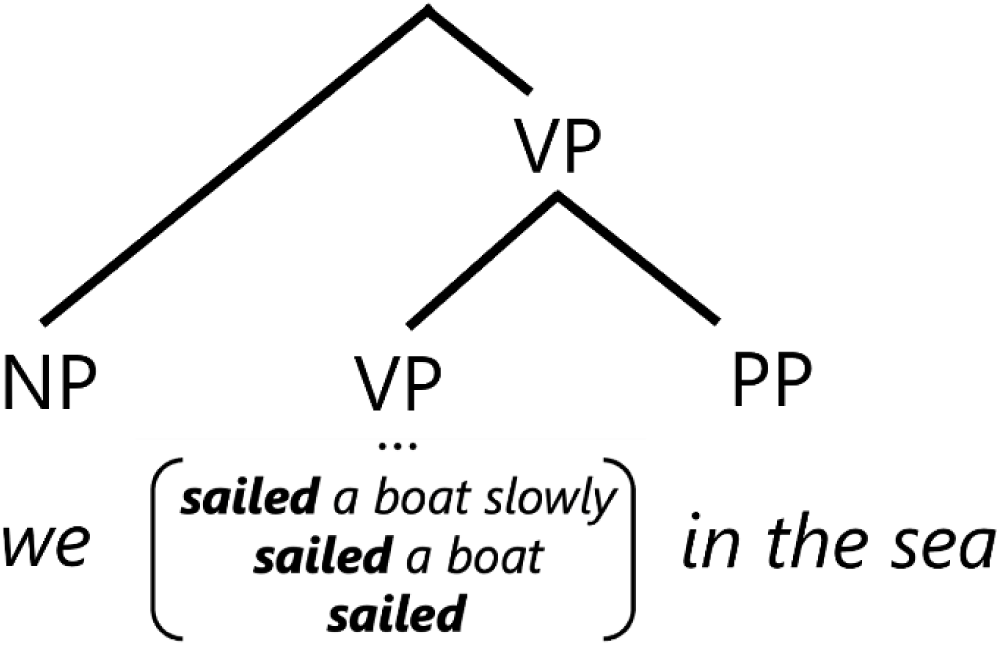
Tree diagram of a sentence with hierarchical headed verb phrases (VPs). Although the verb “sail” forms phrases of different lengths, these phrases fill the same syntactic slot where a VP is required. This VP, in turn, becomes part of a larger phrase.

Probing the representational dynamics of headed phrases is made possible by advances in neural decoding methods that recover from neural activation patterns the underlying cognitive representations, including a word’s PoS (Gwilliams et al., 2023; Murphy et al., 2022). We use model prediction to infer the temporal activation of head information from EEG signals while a phrase is being built. This decoding approach offers not only an interpretable neural readout for structure building, but also a novel application of decoding that goes beyond the accuracy metric (Desbordes et al., 2024; Fyshe et al., 2019; Gwilliams et al., 2023; Murphy et al., 2022).

To test whether the brain builds headed verb phrases (e.g., ***eat*** quickly, where ***eat*** is the verb head), we first train a neural decoder to identify PoS labels (“verb” vs. “adverb”) from EEG signals recorded while participants read sentences in Chinese. Verb activation at each time point in a held-out region that includes a verb-adverb phrase is quantified as the probability that a decoder predicts a “Verb” label, yielding a probability time series (P(v)). The typological properties of Mandarin Chinese enable us to minimally manipulate whether the verb and adverb syntactically compose within the window of analysis. When two words syntactically combine into a phrase, their meanings are also composed (Pylkkänen, 2020) and both processes are found to modulate similar neural signatures (Fedorenko et al., 2016; Parrish & Pylkkänen, 2022). Accordingly, we also orthogonally manipulate the conceptual association between target words to separate conceptual effects.

If the brain builds headed phrases, we expect increased head activation, indexed by P(v), at the phrase-closing point regardless of its proximity to the verbal head. As this computation is structural in nature, we further expect the effect of head reactivation to be independent of conceptual association. Spectral and evoked correlates of these factors are also examined.

## 2. Methods

### 2.1. Participants

35 native speakers of Chinese (14 men, 11 women, mean age = 25.31, SD = 6.27) participated in this study. All participants were right-handed with normal or corrected-to-normal vision. Participants were reimbursed for their time at 20 USD/hour. Written informed consent was obtained prior to the experiment. This study was approved by the Health Sciences and Behavioral Sciences Institutional Review Board at the University of Michigan (Protocol HUM00240913).

### 2.2. Design and Materials

We recorded electroencephalography (EEG) while participants read Mandarin Chinese sentences word-by-word in a rapid serial visual presentation (RSVP) experiment. Experimental materials (Figure 2A) were designed to allow independent data samples for classifier training and model prediction. A neural decoder was first trained on epochs from a non-initial region of the sentence (“training region”) to classify epochs of verbs and adverbs (for decoder training details see section 2.6). The trained decoder was then used to predict PoS labels in a held-out sentence-initial region that has a fixed word order of “V - Particle - Adv” (“prediction region”). Two experimental factors (Phrase closing and conceptual association) were manipulated with a 2 x 2 design in the prediction region at the critical adverb. There was no overlap between training and prediction regions to prevent data leakage in the pipeline.

**Figure 2.**
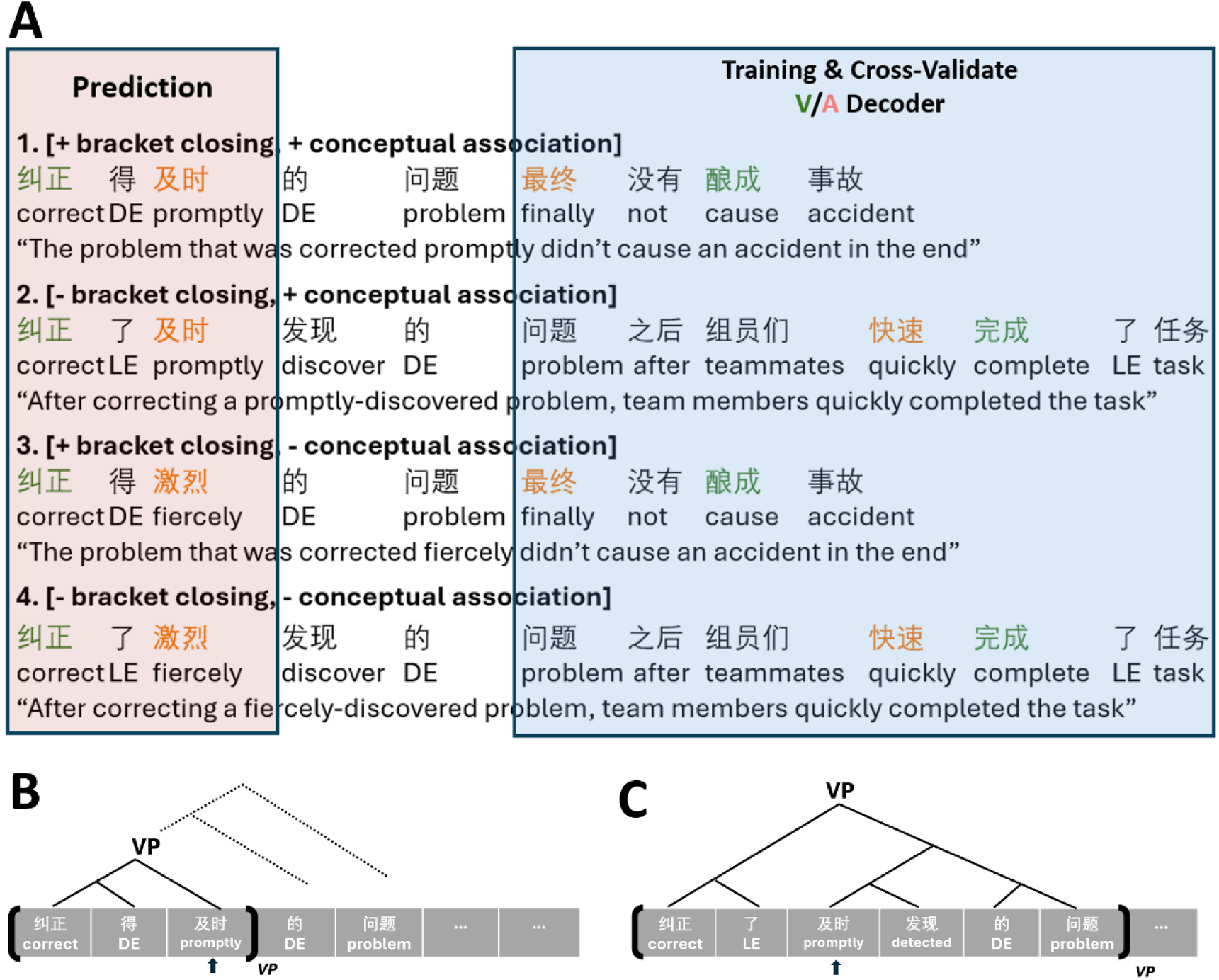
(A) Example of experimental materials from all four conditions illustrates word-segmented sentences in Chinese characters, along with word-by-word translations and English translations. Verbs are in green; adverbs are in orange. (B-C) Syntactic tree diagrams illustrate the phrase closing manipulation. Panel B shows the syntactic structure where the critical adverb (indicated with an arrow) is phrase-final; the VP is formed once the critical adverb “promptly” is presented. Panel C shows the structure where the adverb is not phrase-final: the VP headed by the verb “correct” is closed only after the noun “problem” is presented.

We manipulated two factors in the prediction region that spans the three sentence-initial words: Phrase Closing (with/without a closed verb phrase) and Conceptual Association (high/low association).

Phrase Closing is realized by alternating the Chinese grammatical particle 得/ 了 (DE/LE) which precedes the target adverb. This particle determines whether the initial three words form a grammatical phrase. The particle 得 (DE) marks the target adverb as modifying the sentence-initial verb and closing the VP (e.g., “correct DE promptly” is interpreted as “promptly correct”, Figure 2B); in this condition, the target adverb is phrase final. The particle 了 (LE) marks the phrase as needing a noun-phrase argument; the adverb is interpreted as modifying the following word and, crucially, the VP is not completed until the presentation of the noun (e.g., “correct LE promptly discovered DE problem” is interpreted as “corrected a promptly discovered problem”, Figure 2C). The target adverb thus is phrase-medial, not phrase-final. The syntactic properties of the Chinese DE/LE particles allowed us to manipulate phrase closing without altering any (content) lexical items or word order in the experiment.

Conceptual Association (high/low association) indicates the semantic relatedness between the verb and target adverb in the prediction region. This factor has been shown in previous studies to modulate the same evoked neural signals as phrase closing (Parrish & Pylkkänen, 2022), we thus include it here to test whether phrase decoding can be teased apart from semantic features. High conceptual association materials were created by two native speakers of Mandarin Chinese to form natural sentences. Low conceptual association materials were created by pseudo-randomly shuffling the critical adverb across all sentences. We quantified conceptual association as the cosine similarity between the vector embeddings of a verb-adverb pair in a sentence; Chinese word embeddings come from the Tencent AI Lab (Song et al., 2018). A paired t-test confirms that our manipulation of conceptual association resulted in higher cosine similarity in high association conditions than in low association conditions, *t*(238) = 10.57, *p* < 0.001.

Crossing Phrase Closing (with/without) and Conceptual Association (high/low) resulted in 2 × 2 = 4 conditions. Each condition contained 60 sentences, amounting to a total of 240 sentences. To keep lexical items constant across conditions, a specific verb was always paired with the same adverb (e.g., “correct” with “promptly”) in the “+phrase vs. -phrase” contrast. A lexical difference was inevitable in the high vs. low conceptual association contrast since cosine distance is dependent on the lexical items.

The sentence-final training region forms a natural continuation of the experimental sentences and contains a verb and an adverb per sentence. Verbs and adverbs in the training region were matched in terms of the number of strokes in Chinese characters, word frequency, and word order (i.e., verbs had equal frequency of occurring before and after adverbs).

### 2.3. Procedure

Participants were comfortably seated approximately 100 centimeters from a computer screen in an isolated booth and were instructed to silently read sentences while minimizing movement and eye blinks. Sentences were presented word-by-word at the center of the screen in white text on a grey background using a rapid serial visual presentation (RSVP) protocol. Each sentence began with a fixation cross of 500 ms, followed by individual words for 300 ms and then a 300 ms blank-screen inter-stimulus interval. As an attention check, twenty percent of the trials were followed by a yes/no comprehension question. Participants answered those questions by pressing one of two keys on the keyboard with their left or right index finger. The next trial began immediately after the key press or 500 ms after trials without comprehension questions. Participants completed a short practice session to familiarize themselves with the task. The main session lasted approximately an hour. Participants were given the option of a short break approximately every six minutes during EEG recording.

EEG signals were recorded using an ActiCAP 32-electrode system (Brain Products, Munich, Germany) distributed in a 10-20 layout. The EEG signal was amplified through a BrainAmp ActiCHamp+ DC amplifier, referenced online to a channel placed on the left mastoid, sampled at 500 Hz, and filtered with a passband of 0.1-200 Hz. The impedance of each electrode was kept below 25 kΩ by applying electrolyte gel prior to data recording.

### 2.4. EEG preprocessing

EEG signals were preprocessed and analyzed with the *MNE-python* package (Gramfort et al., 2013; version 1.6.0) in a Python environment. EEG data were re-referenced to the average of the left and right mastoids (TP9, TP10) and then filtered with a bandpass filter to either (1) 1-50 Hz for decoding and time-frequency analyses, or (2) 1-20 Hz for the evoked analysis. Then, the data were epoched from 0.2 s before word onset to 1.2 s post word onset. We used independent component analysis to remove artifacts resulting from eyeblinks and saccades (Jung et al., 2000). Trials with other artifacts such as muscle movement were automatically removed with the *autoreject* Python package (Jas et al., 2017; version 0.4.2). Participants with over 20% trials rejected were excluded from further analysis (epoch rejection rate before exclusion: median = 7.85%, range = 60.69%; after rejection: median = 6.48%, range = 20.14%). Following this criterion, we excluded 5 participants and retained data from 30 participants.

### 2.5. Decoding analysis

To approach our main research question, we trained a PoS decoder to distinguish EEG signals when participants read verbs and adverbs in the sentence-final training region. Then, the trained model was used to generate probabilistic predictions of whether epochs in the sentence-initial prediction region should be classified as having a Verb PoS; we use the term P(v) to denote these temporally resolved probabilistic predictions of verb head information. This serves as our main dependent variable.

#### 2.5.1. Model training

A temporal decoder is a classifier that is trained on spatio-temporal EEG signals via a sliding window, yielding time-resolved estimates. This decoder was trained on verb and adverb epochs from the training region (Figure 3). Epochs were down-sampled to 200 Hz prior to training for faster computing. Similar to Grootswagers et al. (2017), we improved signal-to-noise ratio by (1) averaging two randomly sampled epochs without replacement and (2) using a sliding window of 200 ms and a step size of 50 ms. For each window, a separate logistic regression estimator was trained on data from all channels to predict the correct PoS label associated with the data segment. The estimator was then evaluated with 5-fold cross-validation and summarized in terms of area under the curve of the receiver operating characteristic (“AUC-ROC”). This process is repeated for every 50 ms step through the 1.2-second-long epochs, yielding a time series of cross-validation scores. The sliding-window temporal decoder was trained within-participant such that each participant had a unique temporal decoder that was only trained on epochs from that participant. Cross-validation time series from all participants were then submitted to FDR-corrected paired t-tests to detect time windows of significantly above-chance decoding.

**Figure 3.**
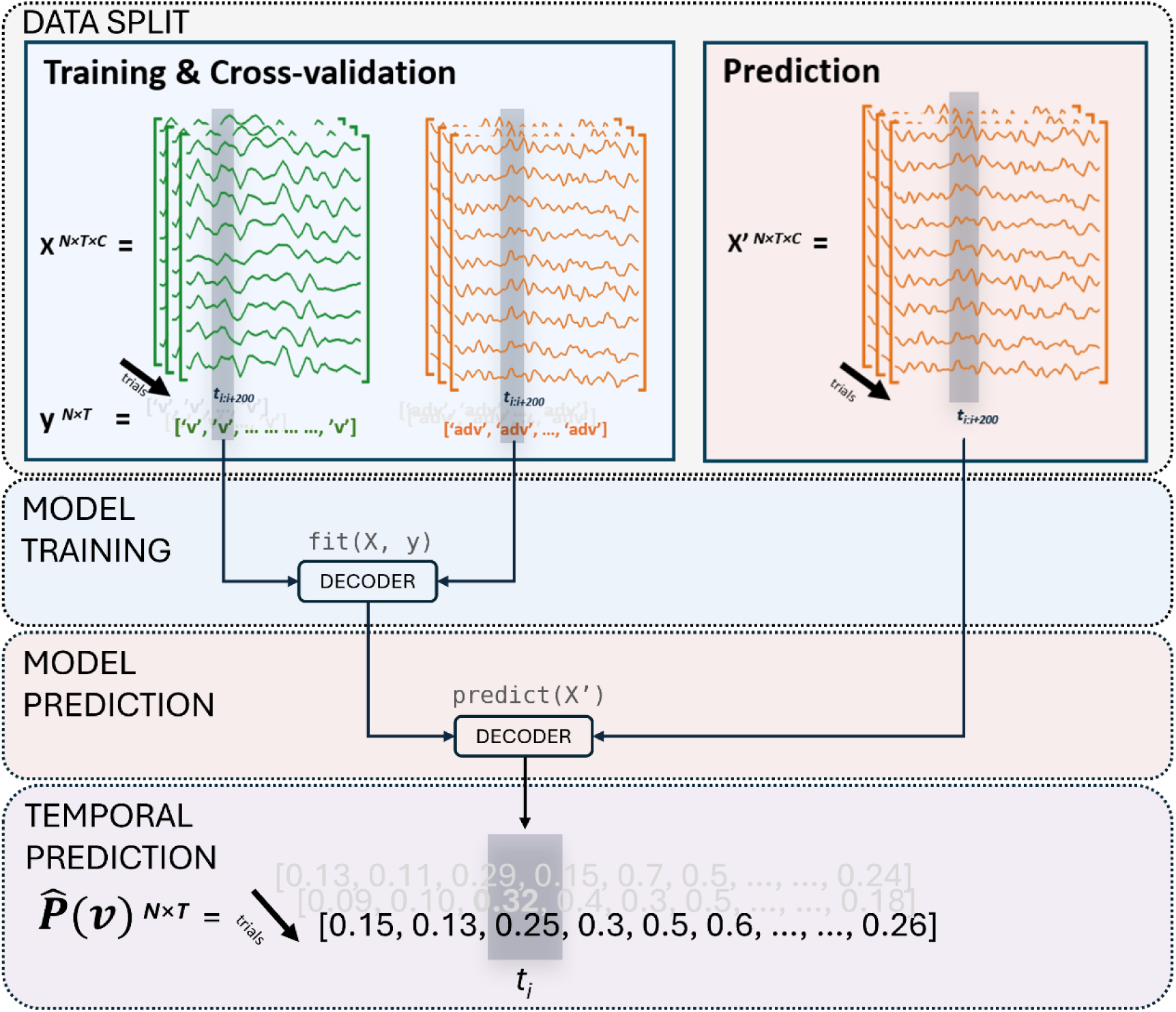
Illustration of the decoding pipeline. A logistic regression estimator was fit for each sliding window to predict the correct PoS label. Training data were 200 ms intervals spanning both verb (green) and adverb (orange) epochs in the sentence-final region. The sliding window moved in 50 ms increments. Trained estimators were used to generate probabilistic verb activation on adverb epochs (word three) in the sentence-initial prediction region, yielding P(v) time series which served as our main dependent variable. To validate our measure, the same testing procedure was performed on verb epochs (word one) in the prediction region, yielding a verb recall time series.

We also implemented two analyses to evaluate the consistency of our decoder in time. First, we extracted topographical decoder patterns (Haufe et al., 2014) over time from trained estimators to examine whether time windows with above-chance decoding had similar scalp distribution of neural activation. Second, we computed a temporal generalization matrix (TGM; King & Dehaene, 2014) to test whether an estimator trained at a specific time point generalizes to an unseen time point in the prediction region (in terms of verb recall, see 2.5.2). Together, these serve to validate the decoding models used to derive our primary measure of interest.

#### 2.5.2. Model prediction

The trained temporal decoder was then used to estimate P(v), the probability that neural signals are classified as verb-like in the sentence-initial prediction region. P(v) was quantified at a given point in time as the probability that a trained decoder returns a “verb” PoS label in a prediction task. As with model training, this step was repeated with a 200 ms sliding window in 50 ms increments. We obtained P(v) time series for all conditions from both verb and adverb epochs in the prediction region. These predictions were derived within-participant too: decoder trained on one participant’s data was only used to derive P(v) for that participant.

P(v) time series obtained **at word three** (i.e., the adverb) in the prediction region served as the critical dependent variable which we interpret in terms of verb activation^1^. If phrase boundaries involve re-activation of the phrasal head, P(v) at the adverb should be higher when that word completes a verb phrase as compared to conditions where the adverb does not complete a verb phrase. We also evaluate P(v) **at the initial verb** in the prediction region; this is equivalent to the model recall metric and this complements the cross-validation check on our analysis. We do not expect any composition-induced difference in P(v) at the initial verb. We constrained our condition-wise comparisons to time windows with above-chance verb recall.

#### 2.5.3. Confound control

Since syntactic EEG effects might extend into a time window that overlaps with the following word, early responses to non-syntactic features at that word might obscure evidence for syntactic effects. We address potential confounds by subtracting responses induced by two non-syntactic factors at the word following the target: number of strokes in Chinese characters (stroke count) and next-word predictability (surprisal). Both factors are believed to modulate early EEG responses (Hauk & Pulvermüller, 2004; Brennan & Hale, 2019).

The subtraction procedure was performed as follows. First, we fit a multiple linear regression model on all epochs with the number of strokes and surprisal as co-regressors, separately for each time-point and each sensor. We then took the residuals of this regression as our de-confounded dependent measure that is orthogonal to any influence of these two factors. Stroke count was derived with the *strokes* Python package (version 0.0.1). Surprisal value at each word was derived from a pretrained Chinese GPT-2 model (Radford et al., 2019; Zhao et al., 2019). Decoder prediction results at the critical adverb are based on de-confounded EEG data.

#### 2.5.4. Significance testing

For statistical evaluation of model evaluation metrics (i.e., cross-validation score, verb recall), we submitted model output at each time point from all participants to a one-sample *t*-test against a null hypothesis of 0.5 (chance level). This procedure was repeated for all time steps. To test if experimental manipulations modulate P(v) at the critical adverb, we ran two-way repeated measures analyses of variance (ANOVA) at each time point to examine the effects of Phrase Closing, Conceptual Association, and their interaction. ANOVAs were constrained to time points with above-chance verb recall, as a meaningful effect on verb activation is contingent on reliable verb decoding.

Multiple comparisons were adjusted with false-discovery rate (FDR) correction (Benjamini & Hochberg, 1995). For the temporal generalization matrix, above-chance performance was detected using a 2-D cluster-based permutation test with 10,000 permutations thresholded at α = 0.05.

### 2.6. Time-locked signal analyses

Following prior work (e.g., Bastiaansen & Hagoort, 2015; Neufeld et al., 2016), we also examined how our manipulation affects time-locked EEG signals, including both evoked responses in the time domain and inter-trial phase coherence (ITPC) in the time-frequency domain.

The evoked EEG response (event-related potential, or ERP) was obtained by averaging the adverb epochs for all four conditions across participants ([-phrase, - association]; [-phrase, +association]; [+phrase, -association]; [+phrase, +association]). To test if there was a significant difference between conditions in the evoked response, we performed a temporal-spatial cluster-based permutation repeated-measures ANOVA with 10,000 permutations (α = 0.05) on all time points and all channels.

Inter-trial phase coherence (ITPC) of the EEG signal was obtained by first transforming original signals into the frequency domain with Morlet wavelets and then computing phase coherence following Cohen (2014):

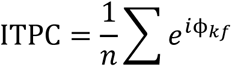

where 𝑒^𝑖ϕ𝑘𝑓^ is the polar representation of the Fourier coefficient with phase angle ϕ at frequency *f* and trial *k* out of *n* trials in total. ITPC is proposed to capture oscillatory neural synchrony of relevance for language understanding at the phrase and sentence levels (Brennan & Martin, 2020; Ding & Simon, 2014; Martin, 2020). The wavelet analysis was applied to linearly spaced frequency bins; wavelets were defined with cycles *c* proportionate to the target frequency band *f*, or *c* = *f* / 2. ITPC of each condition was then represented as the relative signal change from the mean value in a baseline window of -200 – 0 ms. To detect significant time-frequency clusters, we performed a one-sample permutation cluster test (α = 0.05) on ITPC difference at the group level. This test was run on data averaged over a subset of central channels (Cz, C3, C4, CP1, CP2, FC1, FC2) following previous studies using this metric with syntax (Brennan & Martin, 2020; Ding et al., 2017).

## 3. Results

### 3.1. Behavioral comprehension task

Participants achieved an overall accuracy of 87.6% (SD = 8.9%) on comprehension questions that followed 20% of experiment sentences. A repeated-measures ANOVA yielded a significant main effect of Conceptual Association on accuracy such that accuracy was overall higher for sentences with high conceptual association at *w3* (adverb) in the prediction region than those with low conceptual association. *F*(1,28) = 18.91, *p* < 0.001. There is also a significant interaction between Phrase Closing and Conceptual Association, *F*(1,28) = 16.01, *p* < 0.001. High conceptual association led to higher accuracy when there were closed phrases at word 3 (adverb) in the prediction region, *t*(29) = 6.04, *p* < 0.001. Conceptual association did not have a significant impact on accuracy when the phrase was still open in the prediction region, *t*(29) = 0.32, *p* = 0.99. Phrase Closing had no significant main effect on accuracy, *F*(1,28) = 0.27, *p* = 0.60.

### 3.2. Validating PoS decoding in training and prediction regions

A precondition for identifying head activation is reliable decoding of PoS information from EEG data. **In the training region**, a 5-fold cross-validation procedure revealed significant decoding of PoS in 300-350ms and 600-1000ms, *q_max_* = 0.041 (see Figure 4A). **In the testing region**, the trained decoder achieved above-chance verb recall (Figure 4B) in 250-450ms and 550-1000ms, *q_max_* = 0.039. The temporal generalization matrix computed with a training-testing data split (Figure 4C) further revealed that estimators trained in the later 700-1000 ms window consistently generalize to an earlier time window in unseen testing data (100-650ms). The generalization analysis converges with a linear estimator pattern analysis (Fig 4D following Haufe et al., 2014) which suggested similar central-anterior scalp distribution of early and late PoS decoding. Finally, we confirm no significant difference in grammar-external covariates (word frequency, number of strokes in Chinese characters, and GPT-2 surprisal, Figure 4E; pretrained Chinese GPT-2 by Zhao et al., 2019) which may also modulate EEG responses (Brennan & Hale, 2019; Frank et al., 2015; Hauk & Pulvermüller, 2004). We thus consider it unlikely that our models picked up neural activation from features other than PoS.

**Figure 4.**
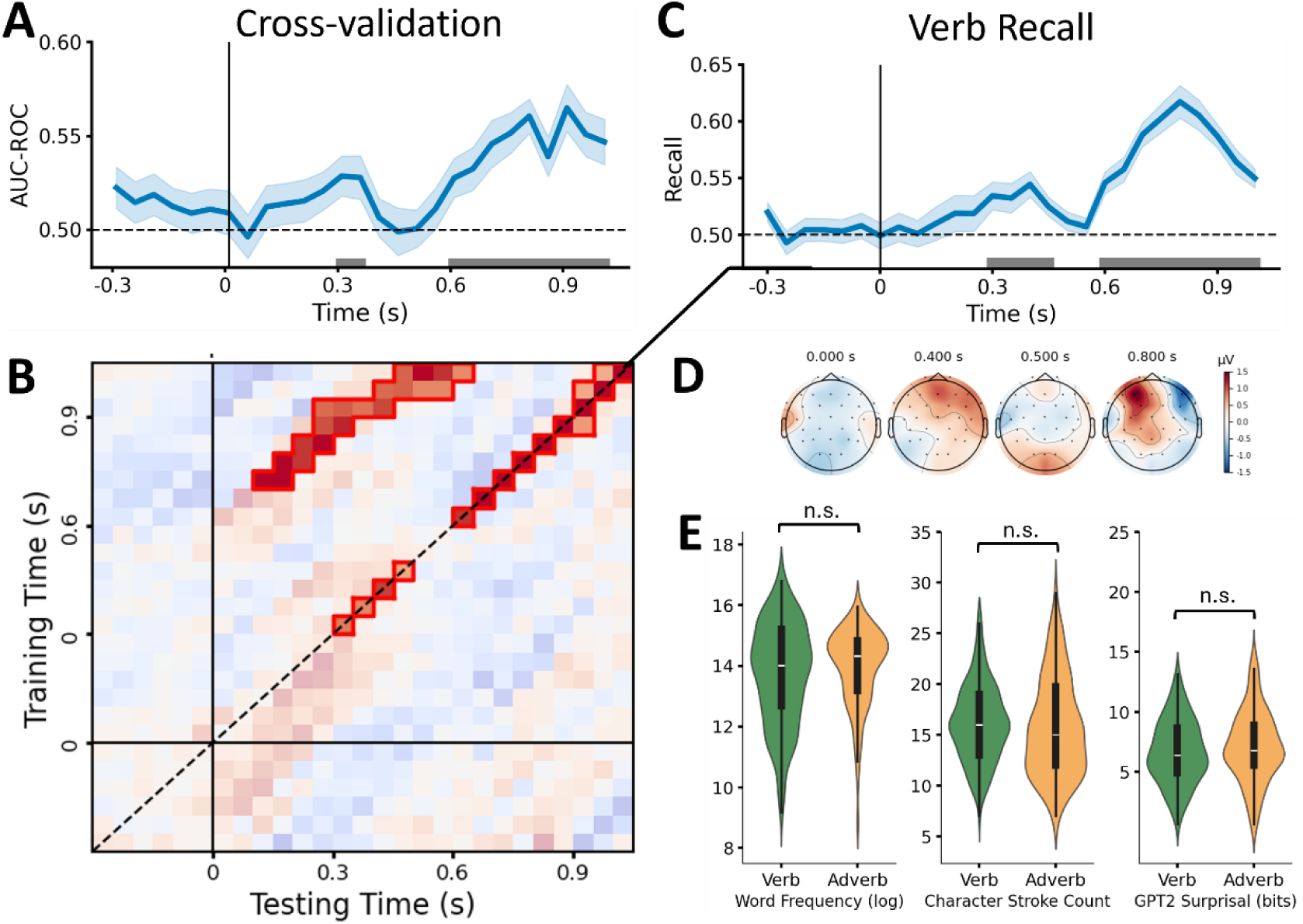
Decoder Performance. (A) Cross-validation score time series from the training region reveal above-chance performance in two windows around 300 ms and 800 ms. Grey bars along the *x*-axis indicate statistically reliable above-chance cross-validation accuracy. Shaded areas indicate ±1 standard error of the mean. (B) Decoder generalizability from training times in the training region to testing times at word 3 (adverb) in the prediction region, quantified in terms of verb recall. Cells with significantly above-chance generalization performance are outlined in red, showing that models trained for the later 800 ms time window reliably classify PoS labels also in the earlier 300 ms window. (C) Verb recall time series validates decoding performance on held-out sentence-initial data; decoding dynamics are similar to that found for cross-validation. (D) Sensor topography patterns from the trained decoder show predominantly frontal involvement in both earlier and later time windows. (E) Statistical distributions of control covariations: log-transformed word frequency, number of characters, GPT2-based surprisal obtained from Chinese verbs and adverbs used in the training region.

In sum, the temporal decoder not only achieved above-chance in the training region, but it also generalizes to unseen data points in a separate sentence-initial testing region and shows consistent spatial patterns across time. We performed model prediction with this decoder to test whether verb activation, operationalized as P(v), is modulated by phrase boundaries.

### 3.3. Grammatical head (verb) activation in the prediction region

The critical dependent variable in this study is the probability that a trained decoder returns a “verb” label during prediction time at the critical adverb; this is P(v). Indeed, we observed an increase in this measure at in a window beginning around 500 ms after onset of the target adverb (Figure 5, top row). Crucially, this increase in P(v) is only observed when the adverb completes a verb phrase; it is not observed when the adverb is phrase-medial. A two-way repeated measures ANOVA revealed a main effect of phrase closing, such that a closed phrase induces higher verb activation 650 to 1000 ms after the critical adverb was presented (*F_min_*(1, 29) = 51.54, *q_max_* = 0.021). This is in line with the hypothesis that the neural representation of heads is reactivated when phrases are constructed. We observed no main effect of Conceptual Association nor an interaction between phrase closing and conceptual association (conceptual association: *F_max_*(1, 29) = 7.67, *q_min_* = 0.135; interaction: *F_max_*(1, 29) = 2.48, *q_min_* = 0.943).

**Figure 5.**
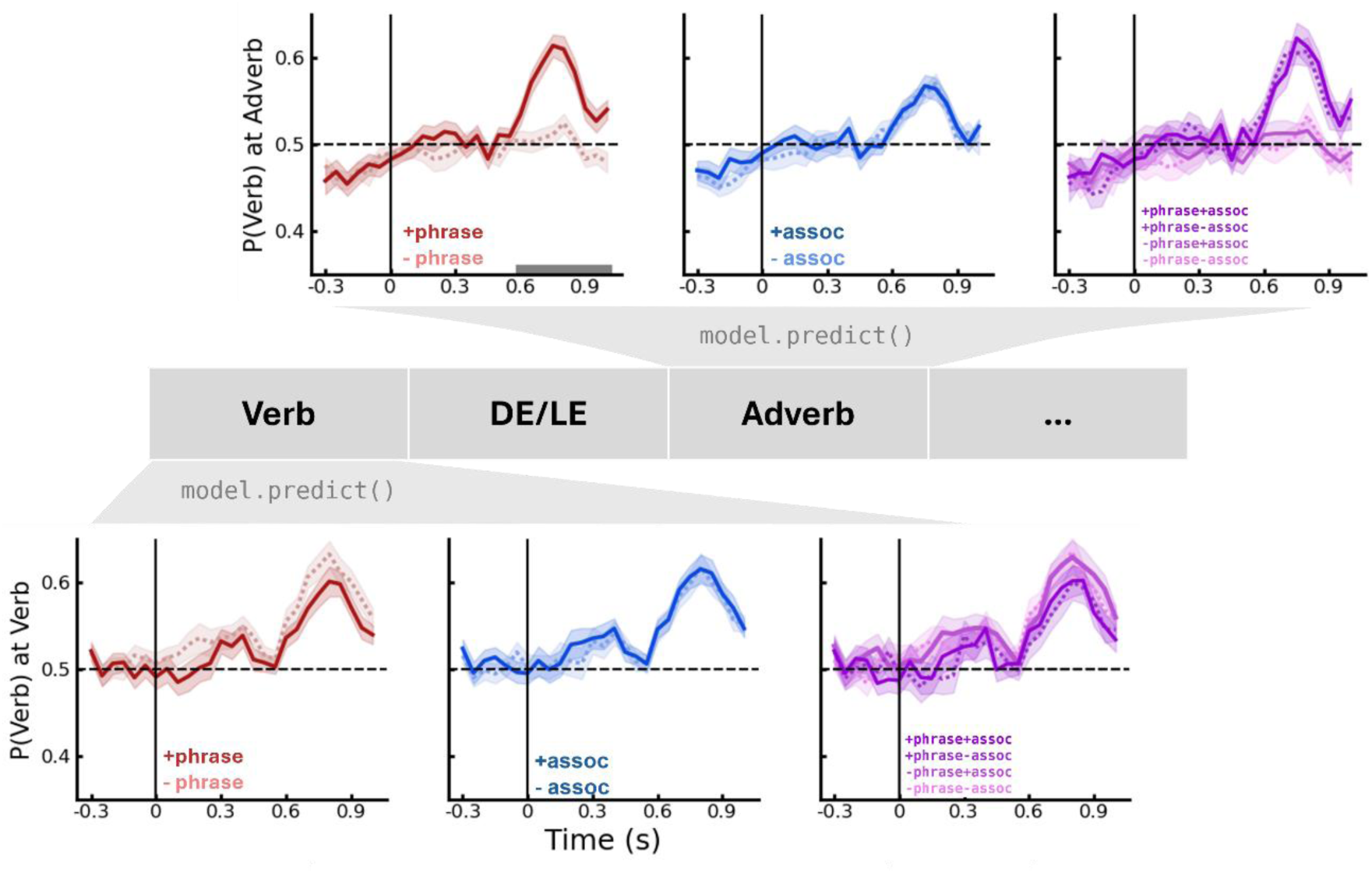
Probabilistic verb activation at the verb (bottom) and adverb (top) in the prediction region. For each row, the panel on the left shows mean P(v) grouped by phrase closing. The panel in the middle represents mean P(v) grouped by conceptual association. The panel on the right represents mean P(v) from all four conditions. Shaded areas indicate ±1 standard error of the mean. Grey bars indicate time windows in which the ANOVA discovered a significant effect. *P*-values used for plotting are FDR-corrected (which are thus *q*-values).

A control ANOVA analysis performed on P(v) obtained at the sentence-initial verb did not find any significant effect of either phrase closing or conceptual association (Fig 5, bottom row; phrase closing: *F_max_*(1, 29) = 6.30, *q_min_* = 0.167; conceptual association: *F_max_*(1, 29) = 1.00, *q_min_* = 0.797).

### 3.4. Control decoding

The results just discussed are based on de-confounded data following the procedure described in section 2.6.3. Although we regressed out non-syntactic effects prior to model prediction, some residual confound in word 4 following the target adverb might still obscure the interpretation of the effect of Phrase closing. To further address this concern, we explicitly trained two decoders to distinguish these covariates *per se* so as to resemble such a “confounded” scenario – we then test the resemblance in terms of performance and decoder weights between the intentionally confounded decoder and the observed results to estimate the potential impact of residual information.

This stroke count decoder was trained following a pipeline similar to that described in section 2.6.1, with the only exception that training epochs were divided along median stroke count (13 strokes) into high and low stroke count. Word type (verbs, adverbs) was balanced in each class. Then, we coded high stroke count epochs as “Adverb” and low stroke count epochs as “Verb”. By doing so, we intentionally confound PoS label with visual complexity as if increased verb activation previously attributed to phrase closing at the critical adverb were indeed due to systematically lower stroke count at the following word. A surprisal decoder was trained similarly, except that we divided the training set along median surprisal.

5-fold cross-validations identified no time windows of significantly above-chance decoding for stroke count (Figure 6A) nor surprisal (Figure 6B). Moreover, at the time point of peak decoding performance (800 ms), the “stroke count confounded” decoder had a positive weight pattern across posterior channels, which is opposite to the central-anterior distribution of positive weight patterns in the actual PoS decoder. The “surprisal confounded” decoder yielded a lateral weight pattern at peak performance (700 ms), which is not comparable with the actual PoS decoder either. This control analysis supports the interpretation of the phrase closing effect above as related to PoS rather than non-syntactic factors.

**Figure 6.**
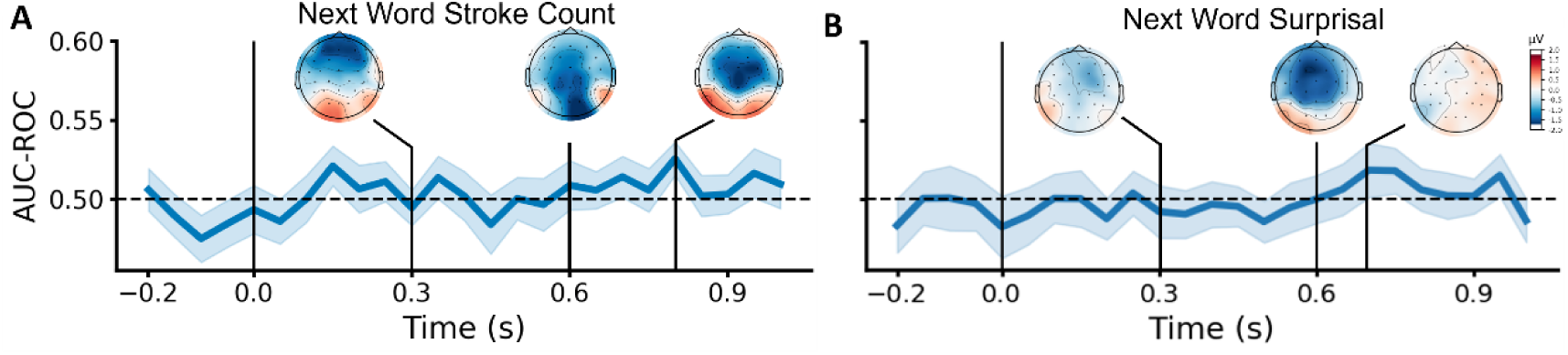
Cross-validation result of control decoders. (A) Control decoder targeting stroke count of word 4. (B) Control decoder targeting surprisal of word 4. Insets show topographical decoder patterns at 300, 600 ms, and the time point with peak performance. No time window reached significance according to FDR-corrected one-sample *t*-tests. Shaded areas indicate ±1 standard error of the mean.

### 3.5. Time-locked analyses

Analysis of evoked EEG response revealed central-anterior negativity spanning from 742 to 894 ms associated with closed phrases (Figure 7A; *p*_cluster_ = 0.002). This effect is still significant when visual complexity and surprisal are regressed out although it was larger when both factors were preserved (Figure 7B; *p*_cluster_ < 0.001). The evoked analysis also revealed an effect of conceptual association, with low association trials eliciting central-anterior negativity in the 354-524 ms time window (Figure 7D; *p_cluster_* = 0.003). We interpret this as an N400 ERP (Lau et al., 2008) and thus a validation of our conceptual manipulation. No interaction between Phrase Closing and Conceptual Association was found.

**Figure 7.**
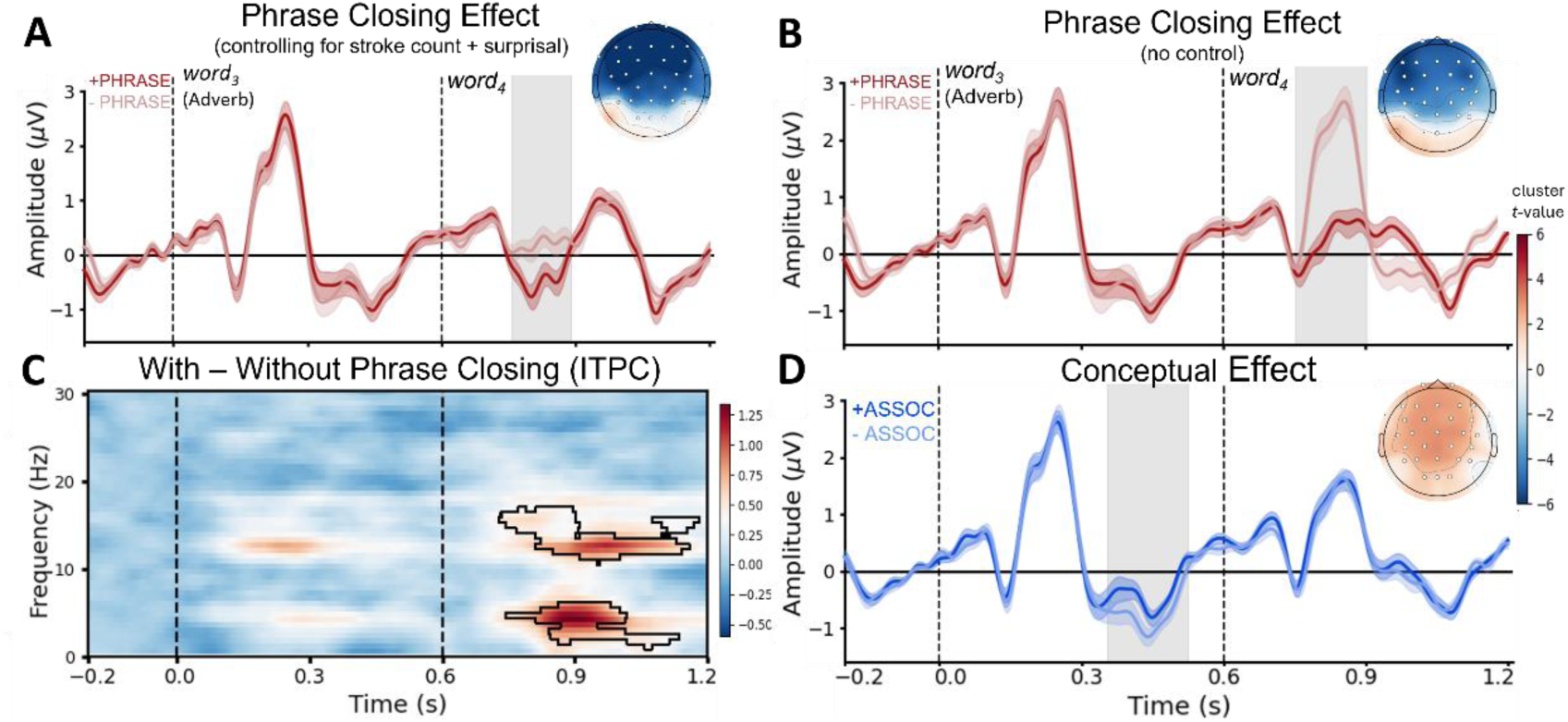
Time-locked results. (A) Evoked response grouped by phrase closing after the effect of stroke count and surprisal was regressed out. (B) Evoked response grouped by phrase closing before the effect of stroke count was regressed out. Grey bands in ERP plots indicate time windows with a significant difference between conditions. Statistical significance is determined by a permutation cluster test thresholded at α = 0.05. Figure insets illustrate the topographical distribution of cluster *t*-values from the permutation test; contrast coding: [1, -1], respective to the order in legends. Channels marked with circles showed a significant effect. Plotted waveforms are the average across sensors that reached significance. (C) Difference in inter-trial phrase coherence induced by phrase closing (after visual confound and surprisal were regressed out). Plotted is the difference in relative ITPC change from a baseline window of -200-0ms. Outlined areas showed a significant effect of phrase closing. (D) Evoked response grouped by conceptual association. Shaded areas indicate ±1 standard error of the mean.

The spectral analysis identified two time-frequency clusters with increased ITPC at closed phrases (Figure 7C). One cluster falls within the delta and low-theta bands (1.1 - 5.9 Hz; *p*_cluster_ = 0.008), spanning 737-1135 ms. The other cluster falls within the beta band (10.7 – 16.8 Hz; *p*_cluster_ = 0.002), spanning 729-1185 ms.

## 4. Discussion

In the current study, we investigate the representational dynamics of phrasal composition with a novel training-prediction neural decoding pipeline. We created a part-of-speech neural decoder that reliably distinguishes brain activity underlying verb and adverb processing. This decoder revealed increased verb activation when the end of a verb phrase is reached. Such a pattern is in line with the account that the neural representation of a phrase shares properties with the grammatical “head” of that phrase (Jackendoff, 1977; Sprouse & Hornstein, 2016). At the verb phrase boundary, we also observed a late negativity in the evoked response and increased inter-trial phase coherence in the delta and beta bands. These findings delineate the dynamics by which phrases are neurally constructed and are consistent with accounts that implicate a role of oscillatory neural activities in linguistic composition (Ding et al., 2016; Martin, 2020). We also showcase a two-step decoding approach to explicitly studying the temporal evolution of neural representations during on-line cognitive processing.

Our decoding approach involves (1) training a PoS decoder and (2) tracking the unfolding of grammatical head (verb) information using model prediction. At the training stage, we achieved significant above-chance decoding of part-of-speech information from the scalp EEG activity recorded in a controlled sentence reading experiment. To date, the neural decoding of grammatical information mostly relies on data with high spatiotemporal resolution (e.g., MEG, ECoG) and/or data recorded while participants read/listened to a very large number of tokens (Desbordes et al., 2024; Gwilliams et al., 2023, 2024; Murphy et al., 2022). By taking a sliding-window approach to improve signal-to-noise ratio (see Grootswagers et al., 2017), we recover PoS information from EEG recordings that have relatively low spatial resolution with a training set of ∼500 tokens per participant. Our decoding configurations are significantly less resource-intensive than previous efforts using EEG, which rely on tens of hours of data from each participant (Murphy et al., 2022).

The trained decoder reliably classifies EEG data into verbs and adverbs in two time windows after target word onset (around 250-500ms and 600-1000ms). Decoder performance is validated on (1) held-out training data from the same sentence region used for training, and (2) unseen data from a different, earlier region in the experimental sentences (see cross-validation and model recall, respectively; Figures 4A, 4B). Model evaluation thus demonstrates that PoS decoding performance can be invariant to word position in a sentence, which is novel in existing decoding studies. The temporal generalizing analysis and decoder pattern analysis further confirmed that the learned representations are spatio-temporally consistent (Figure 4B, 4D).

We hypothesized that headed phrase structure building would involve increased verb activation at the point in the sentence where a VP is closed, even when this point is distal from the verb itself. Model predictions from the temporal PoS decoder showed that this is the case: The model-predicted probability of the “verb” label is higher at the end of a verb phrase, as compared to a sequentially and lexically matched timepoint where no verb phrase closes. We interpret the increased predicted probability for the “verb” label as indexing the reactivation of grammatical head information. In the case of head-initial Chinese verb phrases (VP: V-Adv) used in the current study, increased verb activation was observed 600 ms after the onset of the phrase-closing adverb (or 1800 ms after the verb itself was presented). The phrase closing effect on decoder prediction was accompanied by two time-locked neural patterns: (1) a negative-going event-related component emerging at ∼600 ms, and (2) increased ITPC in the beta and delta bands in approximately 600-1200 ms. None of these effects was modulated by the strength of conceptual association between the verbs and adverbs being composed.

The fundamental role of headed phrase representation in hierarchical language processing is implicated by linguistic theories (Sprouse & Hornstein, 2016) and also behavioral data (Cleland & Pickering, 2003; Coopmans & Schoenmakers, 2020; Frazier & Clifton, 1997; Gibson, 1998; Husain et al., 2014). The present study is the first to explicitly test for these representations in neural signals. Our results are consistent with indirect evidence for the effect of phrasal heads from Zhao et al. (2024). In that study, isochronously presented phrases elicited neural tracking in the delta band and, furthermore, this response was modulated by variability in the position of the head in phrases. Other studies either implicitly assumed such representations (Coopmans et al., 2024), or left underspecified whether hierarchically structured phrases are headed or not (Brennan et al., 2016; Nelson et al., 2017; Shain et al., 2020). The current work provides direct evidence for the neural dynamics of headed phrase representations.

In line with prior work on late electrophysiological activities, we interpret both decoder-derived verb activation and evoked effects in terms of increased syntactic processing load (Hale et al., 2018; Kaan et al., 2000) and phrasal composition (Li et al., 2024). When a phrase is built, the neural encoding of the head is (re)activated as part of the representation of the composed phrase. This process abstracts away from the mere linear order of linguistic input and the conceptual details of constituents. That is, to build a Verb-Adverb phrase, the brain reactivates verb information (as late as ∼1.8 s after the presentation of the verb in the present study), irrespective of how likely the two words conceptually combine. We suggest that such a head (re)activation mechanism is crucial for hierarchical language comprehension: The brain retrieves the information of a previously encountered head (i.e., Verb in Verb-Adverb) and forms a head-centric representation that enables further computations on a larger scale.

Our findings in the time-frequency domain (Figure 7C) point to increased beta- and delta-band activities for headed phrasal structure building. Beta activity is proposed to support memory-based operations such as retrieval (Bastiaansen & Hagoort, 2015; Lewis et al., 2015); delta activity is suggested to aid the extraction of discrete high-level linguistic units from low-level inputs (Ding et al., 2016; Kaufeld et al., 2020; Meyer et al., 2020). These patterns jointly support a mechanism that recruits oscillatory neural activity for headed phrase composition such that delta phase encodes abstract phrase-sized units, beta phase retrieves and binds head information if the head is distant from the phrase-closing point. This mechanism is in line with the proposal that compositional language understanding involves “time-based binding” (Martin, 2020), by which the brain generates endogenous pulses whenever it binds lower-level information (words) to inferred higher-level structures (phrases); temporally aggregated pulses can thus be read out in both frequency and time domains. Our data add detail to this emerging picture in that such higher-level structures involve headed representations.

We did not observe an effect of conceptual association in the decoding analysis or time-locked responses in the time window where phrase closing modulated decoder predictions. This contrasts with some prior work showing that conceptual factors modulate the neural readouts associated with linguistic composition (Fedorenko et al., 2016; Parrish & Pylkkänen, 2022). Since our temporal decoder was trained to distinguish PoS categories that partly dissociate from the mere meaning of words, it is likely that conceptual association does not modulate PoS predictions. Indeed, the fact that PoS decoding was insensitive to conceptual effects supports the validity of our decoder. Regarding time-locked responses, because the current study focused on decoding rather than time-locked analyses, it is by design not directly comparable to the studies that examined the composition of minimal two-word phrases (e.g., Bemis & Pylkkänen, 2011; discussed in more detail below). There is thus indecisive evidence whether conceptual association modulates time-locked readouts that imply syntactic composition.

Two observations in our data merit further discussion. First, the latency of our evoked effect seemingly diverges from prior MEG studies that targeted composition in two-word phrases (see review by Pylkkänen, 2020). These studies suggest that linguistic composition localizes to a time window that centers around 250 ms (Bemis & Pylkkänen, 2011; Flick & Pylkkänen, 2020; Law & Pylkkänen, 2021; Matar et al., 2021); neural activity in this time window is further modulated by the interaction between syntactic and conceptual aspects of phrasal composition (Parrish & Pylkkänen, 2022). We consider three potential causes behind the difference between our data and previous studies: (i) Most studies cited above involved a picture-matching task that requires participants to judge whether a picture matched prior linguistic input or not, whereas in our study participants only read visually presented stimuli with comprehension questions occurring 20% of the time. It is thus likely that our study invokes less multimodal conceptual processing, which results in an effect that localizes to a later time window that is commonly attributed to syntactic processing. (ii) The task we use resembles daily reading more, with longer sentential context, in contrast with minimal two-word phrases. As suggested by Li et al. (2024), variance present in low-level factors potentially delays the composition effect. (iii) The absence of an interaction between syntactic and conceptual factors in our data possibly stemmed from experiment materials. Our [-phrase] manipulation precludes any continuation that is a grammatical Chinese phrase, whereas [-syntactic] phrases used by Parrish & Pylkkänen (2022) involved Adjective-Adjective combinations that are locally grammatical, albeit dispreferred, in English. The presence of potential local grammatical phrases in their [-syntactic] stimuli could introduce milder composition effects that manifest via a semantic main effect or an interaction.

Second, the generalization matrix computed on the train-prediction data split had asymmetric off-diagonal areas (cf. Desbordes et al., 2024; Fyshe et al., 2019): Late estimators generalize well to earlier data increments but early estimators do not generalize to late time points. The earlier time window around 400 ms is commonly linked to processes at the lexical level (Lau et al., 2008) while the later time window is proposed to subserve the processing or re-analysis of grammatical structures (Osterhout & Holcomb, 1992). We thus consider a possible explanation for the asymmetry in generalization performance: Earlier attributes of PoS (e.g., verb: [+concrete]) are less specified than later ones which are required to license fine-grained structure building (e.g, verb: [+concrete, +take animate agent]). This in turn renders a unidirectional entailment from the latter to the former: that a verb is [+concrete, +animate agent] entails it is [+concrete], but a [+concrete] verb does not necessarily entail that it is [+concrete, +animate agent]. Such entailment plausibly translates into the unidirectional generalization (late → early) of PoS decoder performance.

In conclusion, we find evidence for the neural representation of headed phrases with a novel train-prediction decoding pipeline. A neural decoder was trained to decode part-of-speech information from EEG recordings during sentence reading. Subsequent model prediction revealed that head information of a grammatical phrase is reactivated at the phrase closing point, which is accompanied by increased delta- and beta-band activities as well as evoked waveform differences. This work demonstrates an application of neural decoding that uses model prediction, instead of accuracy, to test alternative accounts about a high-level cognitive process in the brain. The results of this study show the role of headed representations in the dynamic neural encoding of phrases and are consistent with accounts of linguistic composition that involve abstract structure building along with the retrieval and binding of head information.

## Acknowledgments

We thank members of the Computational Linguistics Lab and the Psycholinguistics Discussion Group at the University of Michigan for discussions about this project, Cas W. Coopmans for insightful feedback on an earlier version of the manuscript, and Tongyu Lian for inputs on graphic design.

## Author Contributions

J.Z. and J.R.B. designed research; J.Z. and R.G. performed research and analyzed data; J.Z. and J.R.B. wrote the paper.

## Footnotes

1 In the following text, we will refer to this measure as either P(v) or “verb activation” for different purposes.

## References

Bastiaansen, M., & Hagoort, P. (2015). Frequency-based Segregation of Syntactic and Semantic Unification during Online Sentence Level Language Comprehension. Journal of Cognitive Neuroscience, 27(11), 2095–2107. 10.1162/jocn_a_00829

Bemis, D. K., & Pylkkänen, L. (2011). Simple Composition: A Magnetoencephalography Investigation into the Comprehension of Minimal Linguistic Phrases. Journal of Neuroscience, 31(8), 2801–2814. 10.1523/JNEUROSCI.5003-10.2011

Benjamini, Y., & Hochberg, Y. (1995). Controlling the False Discovery Rate: A Practical and Powerful Approach to Multiple Testing. Journal of the Royal Statistical Society. Series B (Methodological), 57(1), 289–300.

Bloomfield, L. (1933). Language. Jason W. Brown Library. https://digitalcommons.rockefeller.edu/jason-brown-library/80

Brennan, J. R., & Hale, J. T. (2019). Hierarchical structure guides rapid linguistic predictions during naturalistic listening. PLOS ONE, 14(1), e0207741. 10.1371/journal.pone.0207741

Brennan, J. R., & Martin, A. E. (2020). Phase synchronization varies systematically with linguistic structure composition. Philosophical Transactions of the Royal Society B: Biological Sciences, 375(1791), 20190305. 10.1098/rstb.2019.0305

Brennan, J. R., Stabler, E. P., Van Wagenen, S. E., Luh, W.-M., & Hale, J. T. (2016). Abstract linguistic structure correlates with temporal activity during naturalistic comprehension. Brain and Language, 157–158, 81–94. 10.1016/j.bandl.2016.04.008

Burroughs, A., Kazanina, N., & Houghton, C. (2021). Grammatical category and the neural processing of phrases. Scientific Reports, 11(1), Article 1. 10.1038/s41598-021-81901-5

Carnie, A. (2021). *Syntax: A Generative Introduction*. John Wiley & Sons.

Chomsky, N. (1957). *Syntactic Structures*. Mouton.

Cleland, A. A., & Pickering, M. J. (2003). The use of lexical and syntactic information in language production: Evidence from the priming of noun-phrase structure. Journal of Memory and Language, 49(2), 214–230. 10.1016/S0749-596X(03)00060-3

Cohen, M. X. (2014). Analyzing Neural Time Series Data: Theory and Practice. MIT Press.

Coopmans, C. W., Hoop, H. de, Tezcan, F., Hagoort, P., & Martin, A. E. (2024). Neural dynamics express syntax in the time domain during natural story listening (p. 2024.03.19.585683). bioRxiv. 10.1101/2024.03.19.585683

Coopmans, C. W., & Schoenmakers, G.-J. (2020). Incremental structure building of preverbal PPs in Dutch. Linguistics in the Netherlands, 37(1), 38–52. 10.1075/avt.00036.coo

Desbordes, T., King, J.-R., & Dehaene, S. (2024). Tracking the neural codes for words and phrases during semantic composition, working-memory storage, and retrieval. Cell Reports, 43(3), 113847. 10.1016/j.celrep.2024.113847

Ding, N., Melloni, L., Yang, A., Wang, Y., Zhang, W., & Poeppel, D. (2017). Characterizing Neural Entrainment to Hierarchical Linguistic Units using Electroencephalography (EEG). Frontiers in Human Neuroscience, 11. 10.3389/fnhum.2017.00481

Ding, N., Melloni, L., Zhang, H., Tian, X., & Poeppel, D. (2016). Cortical tracking of hierarchical linguistic structures in connected speech. Nature Neuroscience, 19(1), Article 1. 10.1038/nn.4186

Ding, N., & Simon, J. Z. (2014). Cortical entrainment to continuous speech: Functional roles and interpretations. Frontiers in Human Neuroscience, 8. https://www.frontiersin.org/article/10.3389/fnhum.2014.00311

Fedorenko, E., Scott, T. L., Brunner, P., Coon, W. G., Pritchett, B., Schalk, G., & Kanwisher, N. (2016). Neural correlate of the construction of sentence meaning. Proceedings of the National Academy of Sciences, 113(41), E6256–E6262. 10.1073/pnas.1612132113

Flick, G., & Pylkkänen, L. (2020). Isolating syntax in natural language: MEG evidence for an early contribution of left posterior temporal cortex. Cortex, 127, 42–57. 10.1016/j.cortex.2020.01.025

Frank, S. L., & Christiansen, M. H. (2018). Hierarchical and sequential processing of language. Language, Cognition and Neuroscience, 33(9), 1213–1218. 10.1080/23273798.2018.1424347

Frank, S. L., Otten, L. J., Galli, G., & Vigliocco, G. (2015). The ERP response to the amount of information conveyed by words in sentences. Brain and Language, 140, 1–11. 10.1016/j.bandl.2014.10.006

Frazier, L., & Clifton, C. (1997). Construal: Overview, Motivation, and Some New Evidence. Journal of Psycholinguistic Research, 26(3), 277–295. 10.1023/A:1025024524133

Friederici, A. D., Chomsky, N., Berwick, R. C., Moro, A., & Bolhuis, J. J. (2017). Language, mind and brain. Nature Human Behaviour, 1(10), 713–722. 10.1038/s41562-017-0184-4

Fyshe, A., Sudre, G., Wehbe, L., Rafidi, N., & Mitchell, T. M. (2019). The lexical semantics of adjective–noun phrases in the human brain. Human Brain Mapping, 40(15), 4457–4469. 10.1002/hbm.24714

Gibson, E. (1998). Linguistic complexity: Locality of syntactic dependencies. Cognition, 68(1), 1–76. 10.1016/S0010-0277(98)00034-1

Gramfort, A., Luessi, M., Larson, E., Engemann, D. A., Strohmeier, D., Brodbeck, C., Goj, R., Jas, M., Brooks, T., Parkkonen, L., & Hämäläinen, M. (2013). MEG and EEG data analysis with MNE-Python. Frontiers in Neuroscience, 7. 10.3389/fnins.2013.00267

Grootswagers, T., Wardle, S. G., & Carlson, T. A. (2017). Decoding Dynamic Brain Patterns from Evoked Responses: A Tutorial on Multivariate Pattern Analysis Applied to Time Series Neuroimaging Data. Journal of Cognitive Neuroscience, 29(4), 677–697. 10.1162/jocn_a_01068

Gwilliams, L., Marantz, A., Poeppel, D., & King, J.-R. (2023). Top-down information shapes lexical processing when listening to continuous speech. Language, Cognition and Neuroscience, 0(0), 1–14. 10.1080/23273798.2023.2171072

Gwilliams, L., Marantz, A., Poeppel, D., & King, J.-R. (2024). Hierarchical dynamic coding coordinates speech comprehension in the brain (p. 2024.04.19.590280). bioRxiv. 10.1101/2024.04.19.590280

Hagoort, P. (2019). The neurobiology of language beyond single-word processing. Science, 366(6461), 55–58. 10.1126/science.aax0289

Hale, J., Dyer, C., Kuncoro, A., & Brennan, J. R. (2018, June 11). Finding Syntax in Human Encephalography with Beam Search. arXiv.Org. https://arxiv.org/abs/1806.04127v1

Haufe, S., Meinecke, F., Görgen, K., Dähne, S., Haynes, J.-D., Blankertz, B., & Bießmann, F. (2014). On the interpretation of weight vectors of linear models in multivariate neuroimaging. NeuroImage, 87, 96–110. 10.1016/j.neuroimage.2013.10.067

Hauk, O., & Pulvermüller, F. (2004). Effects of word length and frequency on the human event-related potential. Clinical Neurophysiology, 115(5), 1090–1103. 10.1016/j.clinph.2003.12.020

Husain, S., Vasishth, S., & Srinivasan, N. (2014). Strong Expectations Cancel Locality Effects: Evidence from Hindi. PLOS ONE, 9(7), e100986. 10.1371/journal.pone.0100986

Jackendoff, R. S. (1977). *X Syntax: A Study of Phrase Structure*. MIT Press.

Jas, M., Engemann, D. A., Bekhti, Y., Raimondo, F., & Gramfort, A. (2017). Autoreject: Automated artifact rejection for MEG and EEG data. NeuroImage, 159, 417–429. 10.1016/j.neuroimage.2017.06.030

Jung, T. P., Makeig, S., Humphries, C., Lee, T. W., McKeown, M. J., Iragui, V., & Sejnowski, T. J. (2000). Removing electroencephalographic artifacts by blind source separation. Psychophysiology, 37(2), 163–178. 10.1111/1469-8986.3720163

Kaan, E., Harris, A., Gibson, E., & Holcomb, P. (2000). The P600 as an index of syntactic integration difficulty. Language and Cognitive Processes, 15(2), 159–201. 10.1080/016909600386084

Kaufeld, G., Bosker, H. R., Oever, S. ten, Alday, P. M., Meyer, A. S., & Martin, A. E. (2020). Linguistic Structure and Meaning Organize Neural Oscillations into a Content-Specific Hierarchy. Journal of Neuroscience, 40(49), 9467–9475. 10.1523/JNEUROSCI.0302-20.2020

King, J.-R., & Dehaene, S. (2014). Characterizing the dynamics of mental representations: The temporal generalization method. Trends in Cognitive Sciences, 18(4), 203–210. 10.1016/j.tics.2014.01.002

Lau, E. F., Phillips, C., & Poeppel, D. (2008). A cortical network for semantics: (De)constructing the N400. Nature Reviews Neuroscience, 9(12), Article 12. 10.1038/nrn2532

Law, R., & Pylkkänen, L. (2021). Lists with and without Syntax: A New Approach to Measuring the Neural Processing of Syntax. Journal of Neuroscience, 41(10), 2186–2196.

Lewis, A. G., Wang, L., & Bastiaansen, M. (2015). Fast oscillatory dynamics during language comprehension: Unification versus maintenance and prediction? Brain and Language, 148, 51–63. 10.1016/j.bandl.2015.01.003

Li, J., Lai, M., & Pylkkänen, L. (2024). Semantic composition in experimental and naturalistic paradigms. Imaging Neuroscience, 2, 1–17. 10.1162/imag_a_00072

Lo, C.-W., Tung, T.-Y., Ke, A. H., & Brennan, J. R. (2022). Hierarchy, Not Lexical Regularity, Modulates Low-Frequency Neural Synchrony During Language Comprehension. Neurobiology of Language, 3(4), 538–555. 10.1162/nol_a_00077

Martin, A. E. (2020). A Compositional Neural Architecture for Language. Journal of Cognitive Neuroscience, 32(8), 1407–1427. 10.1162/jocn_a_01552

Matar, S., Dirani, J., Marantz, A., & Pylkkänen, L. (2021). Left posterior temporal cortex is sensitive to syntax within conceptually matched Arabic expressions. Scientific Reports, 11(1), 7181. 10.1038/s41598-021-86474-x

Meyer, L., Sun, Y., & Martin, A. E. (2020). Synchronous, but not entrained: Exogenous and endogenous cortical rhythms of speech and language processing. Language, Cognition and Neuroscience, 35(9), 1089–1099. 10.1080/23273798.2019.1693050

Murphy, A., Bohnet, B., McDonald, R., & Noppeney, U. (2022). Decoding Part-of-Speech from Human EEG Signals. Proceedings of the 60th Annual Meeting of the Association for Computational Linguistics (Volume 1: Long Papers), 2201–2210. 10.18653/v1/2022.acl-long.156

Nelson, M. J., Karoui, I. E., Giber, K., Yang, X., Cohen, L., Koopman, H., Cash, S. S., Naccache, L., Hale, J. T., Pallier, C., & Dehaene, S. (2017). Neurophysiological dynamics of phrase-structure building during sentence processing. Proceedings of the National Academy of Sciences, 114(18), E3669–E3678. 10.1073/pnas.1701590114

Neufeld, C., Kramer, S. E., Lapinskaya, N., Heffner, C. C., Malko, A., & Lau, E. F. (2016). The Electrophysiology of Basic Phrase Building. PLOS ONE, 11(10), e0158446. 10.1371/journal.pone.0158446

Osterhout, L., & Holcomb, P. J. (1992). Event-related brain potentials elicited by syntactic anomaly. Journal of Memory and Language, 31(6), 785–806. 10.1016/0749-596X(92)90039-Z

Parrish, A., & Pylkkänen, L. (2022). Conceptual Combination in the LATL With and Without Syntactic Composition. Neurobiology of Language, 3(1), 46–66. 10.1162/nol_a_00048

Pylkkänen, L. (2019). The neural basis of combinatory syntax and semantics. Science, 366(6461), 62–66. 10.1126/science.aax0050

Pylkkänen, L. (2020). Neural basis of basic composition: What we have learned from the red–boat studies and their extensions. Philosophical Transactions of the Royal Society B: Biological Sciences, 375(1791), 20190299. 10.1098/rstb.2019.0299

Radford, A., Wu, J., Child, R., Luan, D., Amodei, D., & Sutskever, I. (2019). Language Models are Unsupervised Multitask Learners. https://www.semanticscholar.org/paper/Language-Models-are-Unsupervised-Multitask-Learners-Radford-Wu/9405cc0d6169988371b2755e573cc28650d14dfe

Shain, C., Blank, I. A., van Schijndel, M., Schuler, W., & Fedorenko, E. (2020). fMRI reveals language-specific predictive coding during naturalistic sentence comprehension. Neuropsychologia, 138, 107307. 10.1016/j.neuropsychologia.2019.107307

Song, Y., Shi, S., Li, J., & Zhang, H. (2018). Directional Skip-Gram: Explicitly Distinguishing Left and Right Context for Word Embeddings. In M. Walker, H. Ji, & A. Stent (Eds.), Proceedings of the 2018 Conference of the North American Chapter of the Association for Computational Linguistics: Human Language Technologies*, Volume 2 (Short Papers)* (pp. 175–180). Association for Computational Linguistics. 10.18653/v1/N18-2028

Sprouse, J., & Hornstein, N. (2016). Syntax and the Cognitive Neuroscience of Syntactic Structure Building. In G. Hickok & S. L. Small (Eds.), Neurobiology of Language (pp. 165–174). Academic Press. 10.1016/B978-0-12-407794-2.00014-6

Stanojević, M., Brennan, J. R., Dunagan, D., Steedman, M., & Hale, J. T. (2023). Modeling Structure-Building in the Brain With CCG Parsing and Large Language Models. Cognitive Science, 47(7), e13312. 10.1111/cogs.13312

Zhao, J., Martin, A. E., & Coopmans, C. W. (2024). Structural and sequential regularities modulate phrase-rate neural tracking. Scientific Reports, 14(1), 16603. 10.1038/s41598-024-67153-z

Zhao, Z., Chen, H., Zhang, J., Zhao, X., Liu, T., Lu, W., Chen, X., Deng, H., Ju, Q., & Du, X. (2019). UER: An Open-Source Toolkit for Pre-training Models (arXiv:1909.05658). arXiv. 10.48550/arXiv.1909.05658

